# Development of a nanomedicine approach to improve the therapeutic index of bexarotene

**DOI:** 10.64898/2026.10.05.756654

**Authors:** Sian Rizzo, Carola Pepe, Thibault Albert, Gabriel Braux, Theo Cahart, Carmine Varricchio, Thomas Robertshaw, Timothy R. Hughes, You Zhou, Elaine L Ferguson

## Abstract

Bexarotene (BEX), a retinoid X receptor (RXR) agonist, has demonstrated therapeutic potential in oncology, neurodegenerative diseases and inflammatory conditions, including multiple sclerosis (MS). However, its broader clinical application is limited by poor aqueous solubility, low bioavailability, and dose-limiting adverse effects, most notably hypertriglyceridaemia. In this study, we investigated whether conjugation of BEX to dextrin, a biodegradable polysaccharide, could improve its physicochemical properties and therapeutic index. A dextrin-BEX conjugate was synthesised using succinoylated dextrin (30 mol%) and characterised by gel permeation chromatography (GPC), Fourier-transform infrared (FTIR) spectroscopy, nuclear magnetic resonance (1H-NMR) and dynamic light scattering (DLS). The effects of conjugation on solubility, aggregation behaviour and biological activity were evaluated using *in vitro* cytotoxicity assays in leukaemia, liver and fibroblast cell lines, alongside measurements of intracellular triglyceride levels. The impact on lipid metabolism and toxicity was further assessed in an *in vivo* murine model of hyperlipidaemia. Dextrin conjugation significantly increased the aqueous solubility of BEX (from <0.004 mg/mL to 9.88 mg/mL) and led to the formation of nanoscale aggregates in solution. This was associated with a marked increase in *in vitro* potency, with IC50 values reduced by up to 14-fold across multiple cell lines. However, the increased cytotoxicity was not selective for cancer cells and was also observed in normal fibroblast and hepatic cell models. The dextrin-BEX conjugate did not induce intracellular triglyceride accumulation *in vitro*. *In vivo*, dextrin conjugation did not mitigate BEX-associated lipid dysregulation, while serum triglyceride levels and markers of liver injury remained unchanged. However, although free BEX led to a significant increase in liver weight after 14 days, the dextrin-BEX conjugate did not produce a statistically significant effect. In conclusion, dextrin conjugation markedly improves the solubility and *in vitro* potency of BEX but does not enhance its selectivity or reduce its metabolic toxicity *in vivo*. These findings suggest that while polymer conjugation is an effective strategy for addressing physicochemical limitations, it is insufficient to overcome the inherent dyslipidaemia associated with systemic RXR activation. Future strategies should focus on integrating targeting and controlled release mechanisms, alongside approaches that address the underlying pharmacological drivers of lipid dysregulation.

**Graphical abstract:** 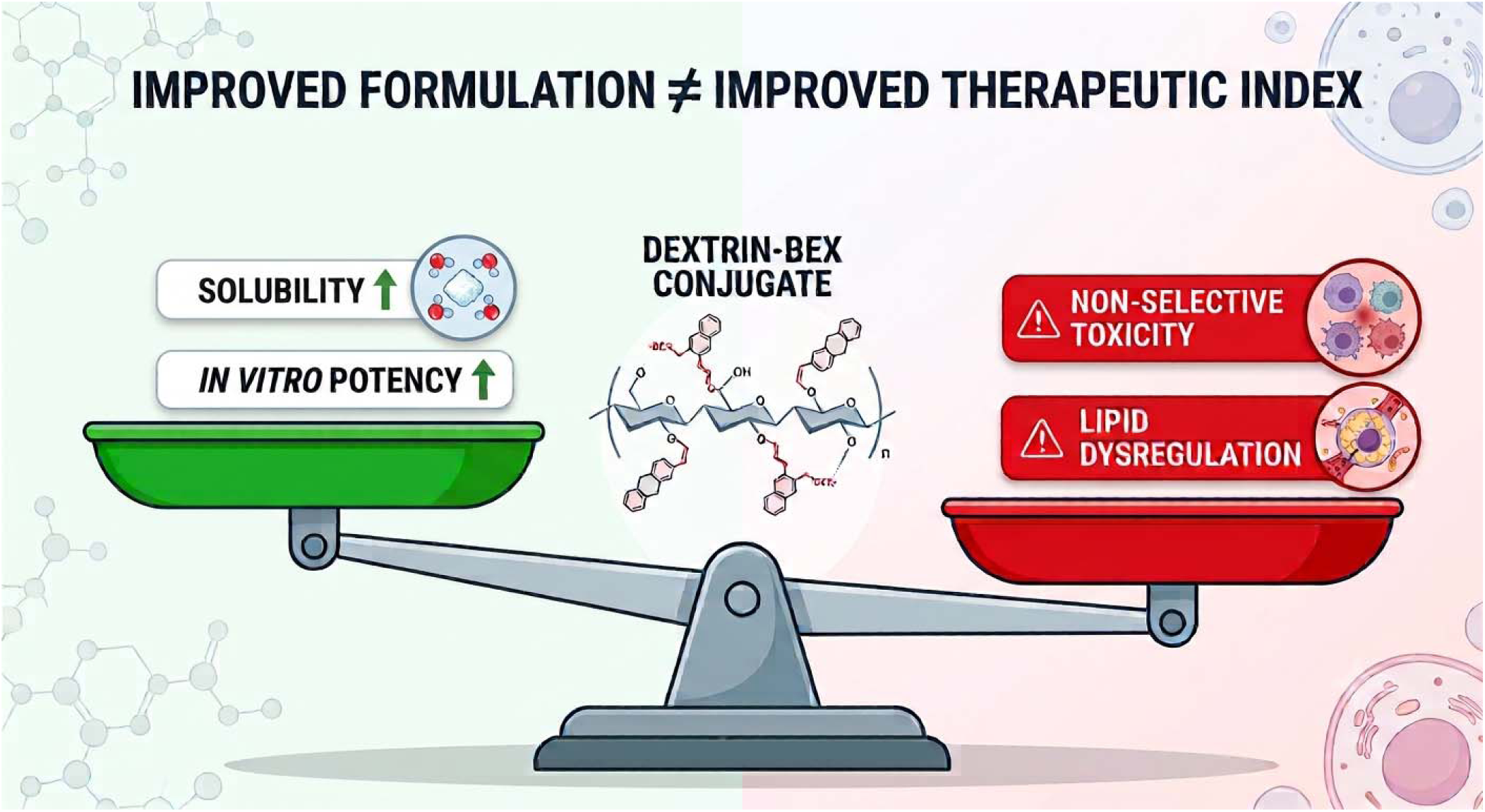

Dextrin conjugation increased bexarotene aqueous solubility and promoted formation of nanoaggregates, leading to enhanced cellular potency. However, the conjugate lacked tumour selectivity and did not mitigate the lipid dysregulation associated with systemic RXR activation in vivo. Graphical abstract generated with the assistance of Google Gemini.

## Introduction

Drug repurposing, or repositioning, is an efficient and cost-effective strategy for expanding the therapeutic applications of established drugs, since their pharmacokinetics and safety profiles are already well characterised.^1^ Bexarotene (BEX), a synthetic retinoid X receptor (RXR) agonist marketed as *Targretin*, was originally developed as an anticancer agent and is approved for the treatment of cutaneous manifestations of refractory advanced-stage cutaneous T-cell lymphoma (CTCL) in adults.^2^ Beyond this indication, BEX has demonstrated therapeutic potential across a range of disease areas, including neurodegenerative disorders, inflammatory conditions and multiple sclerosis (MS), where it has been shown to promote remyelination and improve functional outcomes.^3–7^ Despite these promising applications, the broader clinical use of BEX remains significantly constrained by poor aqueous solubility, low oral bioavailability and dose-limiting toxicities, including hypothyroidism and severe lipid abnormalities.^8, 9^ The intrinsic solubility of BEX is extremely low, necessitating high systemic doses, which can exacerbate adverse effects.^9^ Among these, hypertriglyceridaemia represents the principal barrier to clinical translation, with rates approaching 92% in MS trials and 75% in Alzheimer’s disease studies.^3, 4^ These metabolic effects pose substantial cardiovascular risk and currently prevent the expansion of BEX into non-oncological indications.

To overcome these challenges, numerous drug delivery strategies have been explored to improve BEX solubilisation, pharmacokinetics, and tissue distribution. These include nanocrystals, co-crystals, micelles, liposomes, polymeric nanoparticles and polymer-drug conjugates.^9–17^ Nanocrystal formulations enhance dissolution and bioavailability by reducing particle size, while co-crystals improve solubility and oral absorption, with some systems demonstrating enhanced tissue and brain distribution.^9, 16^ Micellar and lipid-based systems improve aqueous dispersion by encapsulating within hydrophobic domains, although they often require high excipient loads and lack targeting capability.^12, 15^ In parallel, targeted delivery approaches have been developed to improve selectivity and reduce off-target toxicity. Hyaluronic acid (HA)-BEX conjugates, for example, have demonstrated enhanced intracellular localisation and increased anticancer activity with reduced toxicity to normal cells, attributed to receptor-mediated uptake via CD44.^13, 14^ Similarly, ligand-targeted nanoparticle systems, such as folate-functionalised carriers, have shown improved uptake and apoptosis in tumour cells through receptor-mediated internalisation.^10^ However, despite these advances, no delivery strategy to date has demonstrated a clear ability to mitigate BEX-induced dyslipidaemia, suggesting that this adverse effect may be intrinsically linked to RXR activation rather than drug exposure alone.

Polymer-drug conjugation represents a promising strategy to improve the therapeutic index of small molecules by enhancing solubility, prolonging circulation time and modulating biodistribution.^18^ Dextrin is an attractive carrier due to its biocompatibility, biodegradability and long history of clinical use. As a linear polysaccharide, dextrin is enzymatically degraded into non-toxic metabolites (readily degraded by α-amylase to maltose, maltotriose, or maltotetraose,^19^ while chemical modification, such as succinoylation, allows tuning of the degradation rate and drug-release profiles.^20^ Previous studies have shown that dextrin conjugation can reduce drug-related toxicity, as demonstrated by dextrin-colistin conjugates, which exhibited reduced renal accumulation and diminished nephrotoxicity *in vitro* and *in vivo*.^21^

Based on these observations, we hypothesised that dextrin conjugation to BEX could improve its therapeutic index by enhancing solubility and modulating biodistribution, while potentially reducing lipid abnormalities associated with systemic exposure. To test this hypothesis, a dextrin-BEX conjugate was synthesised and evaluated through physicochemical characterisation, *in vitro* cytotoxicity and lipid assays, and an *in vivo* mouse model of hyperlipidaemia. These findings are interpreted within the broader landscape of BEX delivery strategies to determine whether polymer conjugation can address the key translational limitations of this drug.

## Materials and methods

### Materials

Dextrin (Mw 51,000 g/mol, DP = 315) was from ML Laboratories (Liverpool, UK). Bexarotene, dimethyl sulfoxide (DMSO), human insulin solution, ammonium acetate, Δ^9^-tetrahydrocannabinol solution and staurosporine were purchased from Merck Life Science UK Limited, Gillingham, UK). Disodium hydrogen phosphate, potassium dihydrogen phosphate, potassium chloride, succinic anhydride, 4-dimethylaminopyridine (DMAP), acetonitrile (ACN), glacial acetic acid, BCA protein assay kit, 1-ethyl-3-(3-(dimethylamino)propyl carbodiimide hydrochloride) (EDC), sodium chloride, 0.05% w/v trypsin-0.53 mM EDTA, Iscove’s Modified Dulbecco’s Medium (IMDM) with GlutaMAX^TM^, Roswell Park Memorial Institute (RPMI) 1640 medium with GlutaMAX^TM^, Dubelcco’s Modified Eagle Medium (DMEM) with high glucose (4.5 g L^−1^) and GlutaMAX^TM^, William’s medium E, MEM amino acids supplement and foetal bovine serum (FBS) were obtained from ThermoFisher Scientific (Loughborough, UK). Recombinant human granulocyte-monocyte colony-stimulating factor (rhGM-CSF) was from Peprotech (London, UK). Pullulan gel filtration standards (M_w_ = 180−788,000 g/mol) were purchased from Polymer Laboratories (Church Stretton, UK). Unless otherwise stated, all chemicals were of analytical grade and used as received. All solvents were of general reagent grade (unless stated) and were from Fisher Scientific (Loughborough, UK).

### Cell Culture

The cell lines MV-4-11 (RRID:CVCL_0064), TF-1 (RRID:CVCL_0559) and HEpG2 (RRID:CVCL_0027) were obtained from ATCC (Manassas, USA). THP-1 (RRID:CVCL_0006) cells were obtained from ECACC (UK Health Security Agency). Immortalised human hepatocytes (IHH) (RRID:CVCL_8278)^22^ and Huh7 (RRID:CVCL_0336)^23^ cells were provided by Dr You Zhou (Cardiff University). hTERT-immortalised fibroblast cells from patient 2 (nIh)^24, 25^ were provided by Prof Phil Stephens (Cardiff University). Characteristics and culture conditions of all cell lines are summarised in Table S1.

Cells were maintained in log-phase growth at 37°C in a humidified atmosphere containing 5% CO_₂_ and cultured in their respective media. Cells were screened for mycoplasma contamination upon thawing and at monthly intervals thereafter. Subculture was performed using 0.05% w/v trypsin-0.53 mM EDTA.

### Animals

All animal experiments were conducted in accordance with the United Kingdom Animals (Scientific Procedures) Act 1986. Procedures were reviewed and approved by the Animal Welfare and Ethical Review Body under the establishment licence held by Cardiff University and authorised by the UK Home Office. Female C57BL/6J mice (8 weeks old) were purchased from Charles River Laboratories (Bristol, UK). Animals were acclimatised for 7 days prior to experimentation and maintained under standard laboratory conditions in accordance with institutional guidelines.

### Synthesis and characterisation of dextrin-BEX conjugate

First, succinoylated dextrin (30 mol%) was synthesised and characterised as previously described,^26^ then conjugated to BEX (Figure 1). Briefly, BEX (100 mg, 0.287 mmol), EDC (110 mg, 0.574 mmol) and DMAP (35 mg, 0.287 mmol) were dissolved in anhydrous DMSO (5 mL) under stirring in a 50 mL round-bottomed flask for 15 min. A solution of succinoylated dextrin (465 mg, corresponding to 2.87 mmol monomer units) in anhydrous DMSO (5 mL) was then added, and the reaction mixture was stirred for 24 h at 50°C. The resulting conjugate was purified using Sephadex G-25 PD-10 desalting columns equilibrated in Milli-Q water. The product was further concentrated using Vivaspin centrifugal filter tubes (MWCO 3 kDa) by centrifugation (15 min at 4500 rpm) followed by washing with Milli-Q water for three cycles. The final conjugate was lyophilised and stored at −20°C.

**Figure 1.**
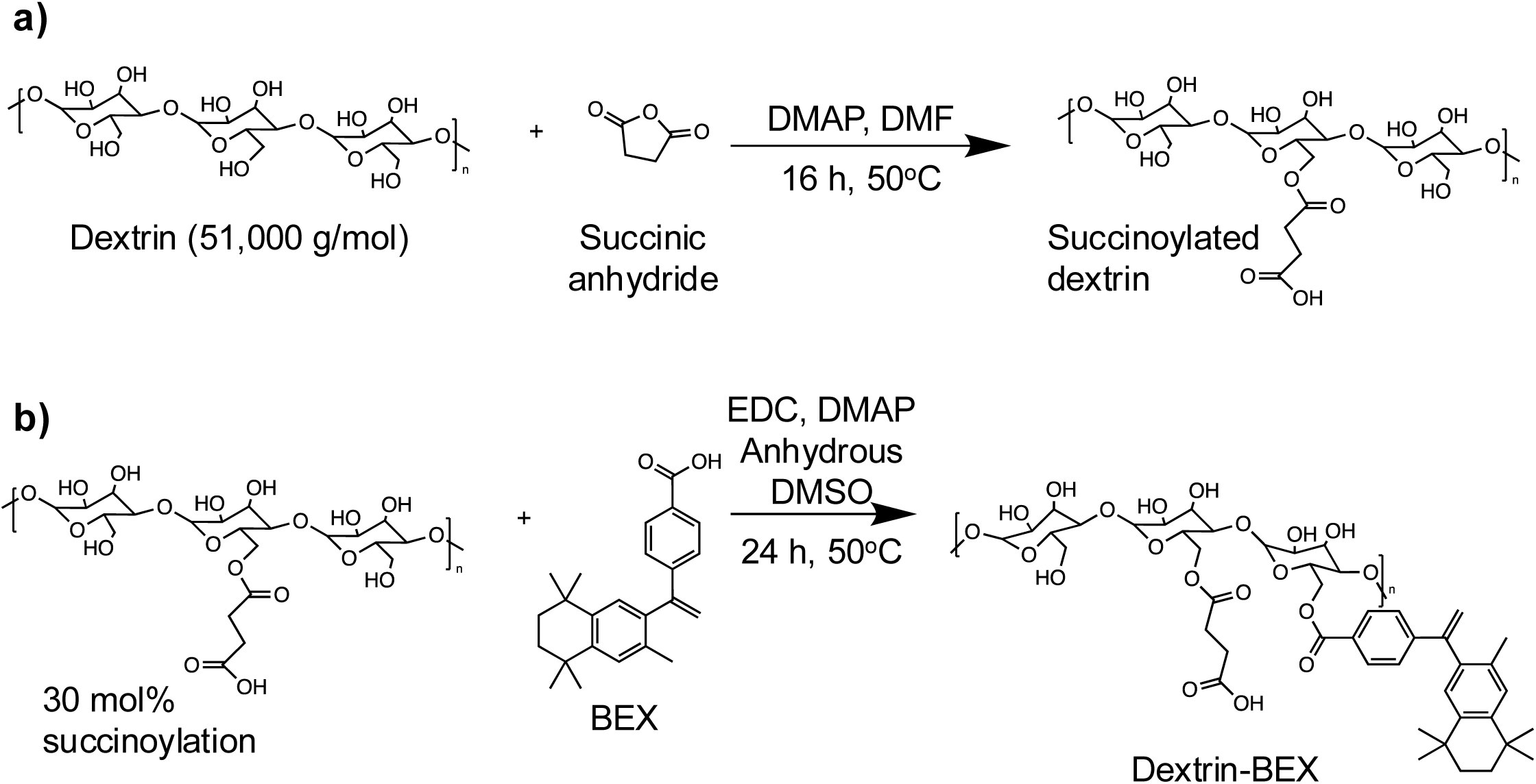
Scheme for synthesis of a) succinoylated dextrin and b) dextrin-bexarotene (BEX) conjugate.

### Physicochemical characterisation

Gel permeation chromatography (GPC) was used to assess molecular weight, purity and polydispersity relative to pullulan standards. The GPC system consisted of TSK G4000PWXL and G3000PWXL columns (Polymer Laboratories, Church Stretton, UK) connected in series to a Jasco PU-1580 HPLC pump, with phosphate-buffered saline (PBS) as the eluent at a flow rate of 1 mL/min. Samples (3 mg/mL, 200 µL) were injected into a 100 µL loop, and detection was performed using a Gilson 153 differential refractometer (Middleton, WI, USA) and a Jasco MD-2010 multi-wavelength UV detector (260 and 280 nm). Detectors were connected to a Polymer Laboratories PL DataStream (Church Stretton, UK) and data were analysed using Cirrus GPC software (version 3.2, 2006) from Polymer Laboratories (Church Stretton, UK).

BEX content of the conjugate was determined by UV spectroscopy at 260 nm using a calibration curve prepared from BEX standards (3.9-125 µg/mL in PBS).

Fourier-transform infrared (FTIR) spectroscopy and proton nuclear magnetic resonance (1H-NMR) were used to confirm the formation of ester linkages and verify the chemical structure.

DLS measurements were performed using a DynaPro Plate Reader III (Wyatt Technology Europe GmbH, Dernbach, Germany). Stock solutions of BEX and dextrin-BEX conjugate were prepared in DMSO (50 mg/mL) and serially diluted (two-fold) in Milli-Q water to obtain concentrations equivalent to 0.07-17.9 µM BEX. Aliquots (150 µL) were transferred to black, clear-bottom 96-well plates and centrifuged (5 min at 1000 *g*) prior to analysis. Samples were measured at 37°C using 10 acquisitions, each lasting 10 s, in auto-attenuation mode. Each sample was measured in duplicate, with two independent experiments performed. Particle size (hydrodynamic radius) and scattering intensity were determined using DYNAMICS^®^ software 8.2 (Wyatt Technology Corporation, Santa Barbara, CA, USA). The onset of aggregation was estimated by plotting scattering intensity (kcps) versus concentration, analogous to determining a critical aggregation concentration.

### Water solubility

The water solubility of BEX and the dextrin-BEX conjugate was determined by preparing saturated solutions. Excess amounts of BEX (20 mg) or dextrin-BEX conjugate (50 mg) were added to 1 mL Milli-Q water and stirred for 1 h at 20°C. Suspensions were centrifuged (8000 g for 10 min), and supernatants were collected and diluted (BEX 1:10, dextrin-BEX 1:200) prior to analysis. UV absorbance was measured at 260 nm, and concentrations were calculated using the BEX calibration curve. Samples were prepared in triplicate.

### Evaluation of *in vitro* cytotoxicity

Multiplexed assays (Promega, WI, USA) were used to assess metabolic activity (CellTiter Blue), apoptosis (Caspase-Glo® 3/7), and necrosis (CytoTox-One®) following treatment, as previously described.^21, 27^ Cells were exposed to BEX, dextrin-BEX conjugate, succinoylated dextrin or vehicle alone for up to 72 h. Additional controls included unconjugated BEX, combined with succinoylated dextrin at equivalent concentrations as used in dextrin-BEX conjugates, staurosporine (10 μM: THP-1, MV-4-11; 1 μM: TF-1, HepG2, Huh7, IHH) as a positive control for apoptosis, and Triton X-100 (100 μM) as a positive control for necrosis. Cell-free (complete media), untreated cells (cells + complete media) and vehicle control (cells + complete media + PBS/DMSO) were also included.

Cells were plated and randomly assigned to treatment groups, then treatments added according to a planned layout. Cells were prepared with 3 replicates per plate (10,000 cells/well in 100 μL, n=3) and each experiment was performed at least twice. Plates were read using a BMG FLUOstar Omega plate reader (BMG Labtech, Aylesbury, UK).

### *In vitro* analysis of triglyceride production

Intracellular triglyceride levels were measured in HepG2, Huh7 and IHH cells using a Triglyceride-Glo assay kit (Promega, Southampton, UK). Cells were treated with test compounds for up to 48 h, with vehicle-treated cells (cells + complete media + PBS/ DMSO) serving as controls. Cells were plated and randomly assigned to treatment groups, then treatments added according to a planned layout. Cells were seeded at 10,000 cells per well (100 µL) in triplicate, and experiments were performed twice. Plates were read using a BMG FLUOstar Omega plate reader (BMG Labtech, Aylesbury, UK).

### *In vivo* hyperlipidaemia model

Mice were randomly assigned to four groups (n=6 animals each): BEX, dextrin-BEX conjugate, succinoylated dextrin, and saline as a control. Treatments were given intraperitoneally every 24 hours for 14 days at a dose of 7.5 mg BEX base per kg per day (total 105 mg/kg). This dosage was selected based on prior research: oral BEX at 30-40 mg/kg for up to 3 weeks notably increased triglyceride and cholesterol levels,^28, 29^ and 5 mg/kg BEX via IP route for 14 days showed no toxicity.^30^ Since prior *in vitro* studies indicated approximately a 1-log increase in toxicity with dextrin conjugation, the dose was kept at 7.5 mg/kg to prevent unforeseen toxicity. To reduce bias, animals were identified with coded ear tags. Body weight was measured daily. Blood samples were taken before treatment and on days 1 and 7, then stored as serum at −20°C.

Twelve hours after the final dose, mice were euthanised using CO_₂_, followed by cardiac puncture. Blood samples were collected, and serum was separated by centrifugation (2000 g for 10 min) and stored at −80°C. Livers were perfused with PBS, harvested and snap-frozen.

Liver tissue was homogenised in PBS using gentleMACS M tubes (Miltenyi Biotec Ltd, Surrey, UK) and a gentleMACS dissociator using the standard protein program. Total protein content was determined using a BCA assay.

### Biochemical analysis

Serum aspartate aminotransferase (AST) activity was measured using a commercial assay kit (Merck Life Science UK Limited, Gillingham, UK). Serum and liver triglyceride levels were determined using a colourimetric assay (Cayman Chemical, MI, USA).

Liver cytokine levels (IL-6 and TNFα) were measured using DuoSet ELISA kits (R&D Systems, MN, USA). Samples (n = 6 per group) were analysed in duplicate and normalised to control values.

### Measurement of BEX tissue levels

BEX concentrations in serum and liver homogenates were measured using HPLC-UV using a method adapted from Lee et al. 2017.^31^ Briefly, HPLC-UV analysis was performed using a Dionex ICS-3000 ion chromatography system (Thermo Scientific, Gloucester, UK) equipped with a Dionex AS autosampler maintained at 4°C. Data were collected and processed using Chromeleon 6.8 software. Chromatographic separation was achieved using a Waters Atlantis C18 column (5 μm particle size, 2.1 × 150 mm; Milford, MA, USA) connected to a SecurityGuard guard column (3 μm particle size, 2 × 4 mm; Phenomenex, Macclesfield, UK), housed in a column oven at 40°C. The mobile phase, consisting of acetonitrile (ACN) and ammonium acetate buffer (10 mM, pH adjusted to 4.1 with glacial acetic acid) in a 75:25 (v/v) ratio, was delivered under isocratic conditions at a flow rate of 0.2 mL/min, with analytes detected at 259 nm. Samples were prepared by protein precipitation with ACN followed by liquid-liquid extraction using n-hexane:ethyl acetate (10:1, v/v), according to the method described by Lee et al.^31^ The injection volume was 50 μL, and the total run time was 25 min. Δ9-tetrahydrocannabinol (250 ng/mL) was used as the internal standard (IS). The retention times for BEX and IS were ∼10.8 and 14.8 min, respectively. Quantification of BEX was performed using a calibration curve constructed from a series of standards (0, 312.5-10,000 ng/mL) prepared in ACN containing IS and subjected to the same sample preparation procedure. Samples (n = 6 per treatment) were analysed in duplicate.

### Statistical analysis

Statistical analysis was performed using GraphPad Prism (version 9.3.1, San Diego, CA, USA). Data are presented as mean ± SEM. Technical replicates were excluded only when values were at least 2-fold above or below the mean of the remaining replicates. Data values outside a biologically plausible range were excluded from the analysis (e.g., negative absorbance values). One-way analysis of variance (ANOVA) followed by Dunnett’s post hoc test was used for multiple comparisons unless otherwise stated. Statistical significance is indicated by *, where * *p* < 0.05, ** *p* < 0.01, *** *p* < 0.001 and **** *p* < 0.0001.

## RESULTS

### Synthesis and characterisation

Two batches of dextrin-BEX conjugates were prepared for use in this study; their characteristics are summarised in Table 1.

**Table 1.**
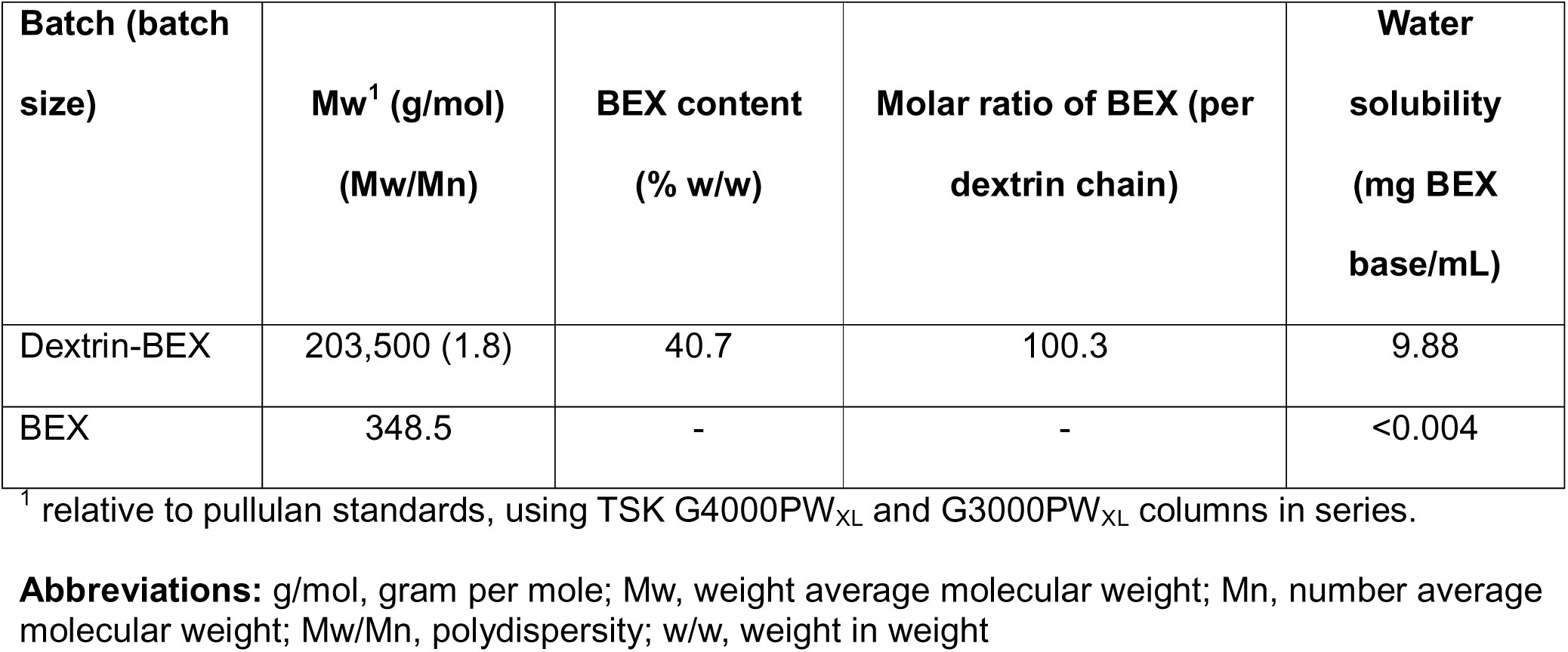
Characteristics of dextrin-BEX conjugate batches.

| Batch (batch size) | Mw<sup>1</sup> (g/mol)<br>(Mw/Mn) | BEX content<br>(% w/w) | Molar ratio of BEX (per dextrin chain) | Water solubility<br>(mg BEX base/mL) |
| --- | --- | --- | --- | --- |
| Dextrin-BEX | 203,500 (1.8) | 40.7 | 100.3 | 9.88 |
| BEX | 348.5 | - | - | <0.004 |

<sup>1</sup> relative to pullulan standards, using TSK G4000PW<sub>XL</sub> and G3000PW<sub>XL</sub> columns in series.
**Abbreviations:** g/mol, gram per mole; Mw, weight average molecular weight; Mn, number average molecular weight; Mw/Mn, polydispersity; w/w, weight in weight

GPC confirmed the presence of high molecular weight conjugates, while FTIR and ^1^H-NMR confirmed the formation of ester bonds between succinoylated dextrin and BEX. FTIR and ^1^H-NMR of free BEX and dextrin-BEX conjugates are shown in Figure 2 and S1. DLS analysis showed that dextrin-BEX conjugates had a higher propensity to aggregate than the free drug, forming aggregates above 0.283 μM (Figure 2d).

**Figure 2.**
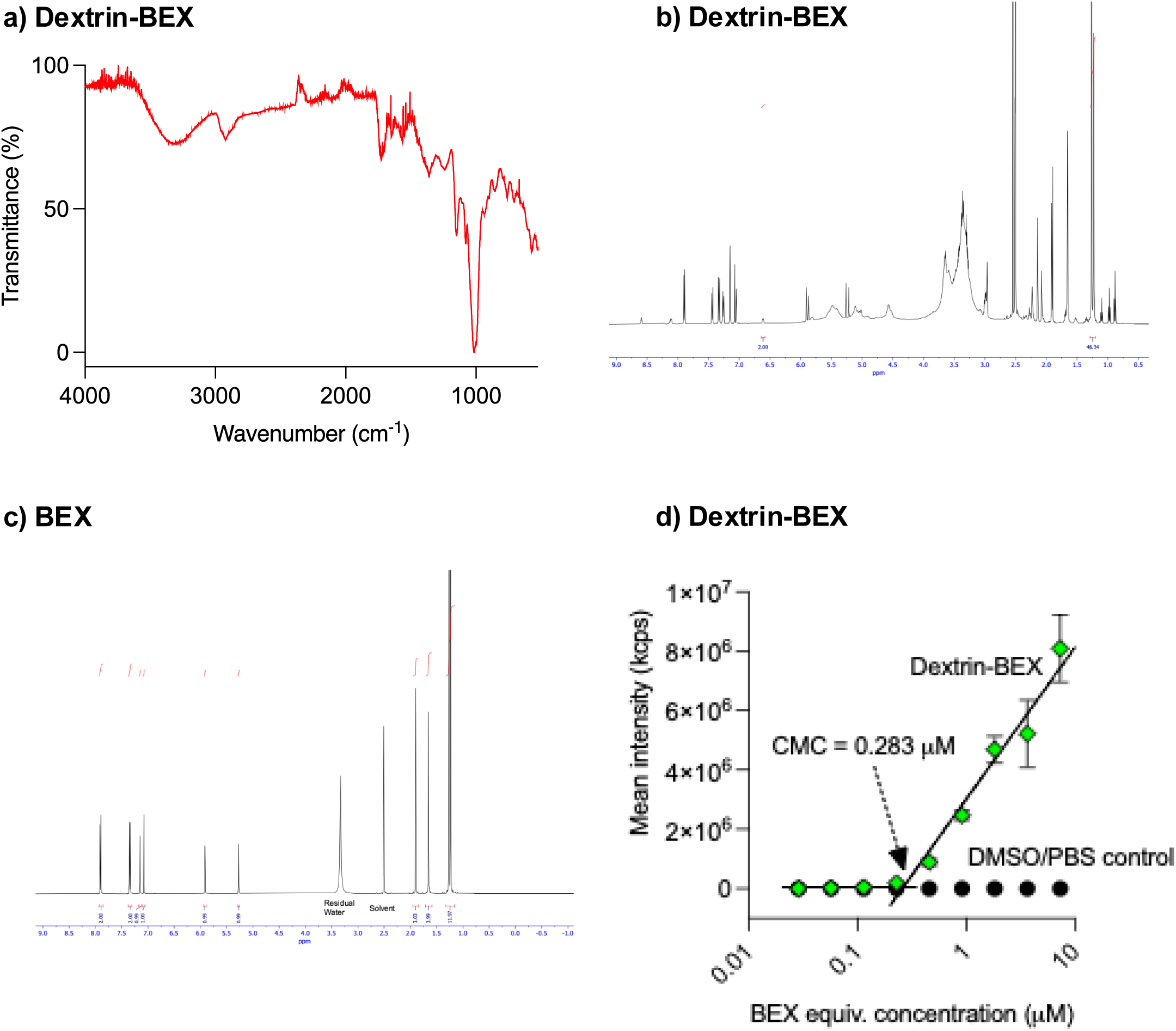
Physicochemical characterisation of DexBEX conjugates. Panels show a) DLS, b) FTIR, and c) ^1^H-NMR of dextrin-BEX. Panel d) shows ^1^H-NMR of BEX.

BEX caused a dose-dependent decrease in cell viability of all leukaemia cell lines tested (Figure 3a, S2). Conjugation of succinoylated dextrin increased BEX’s cytotoxicity (by up to 14-fold), resulting in a significantly reduced IC50 value (p<0.0001, Table 2), and enhanced cell death via necrosis (Figure S3) and caspase 3/7-dependent apoptosis (Figure 3b, S4). Prolonged incubation of cells (up to 72 h) with the treatments caused more pronounced effects on cell viability (data not shown). In contrast, succinoylated dextrin, at equivalent doses as found in dextrin-BEX conjugate preparations, had no effect on cell viability, even at the highest concentration tested.

**Figure 3.**
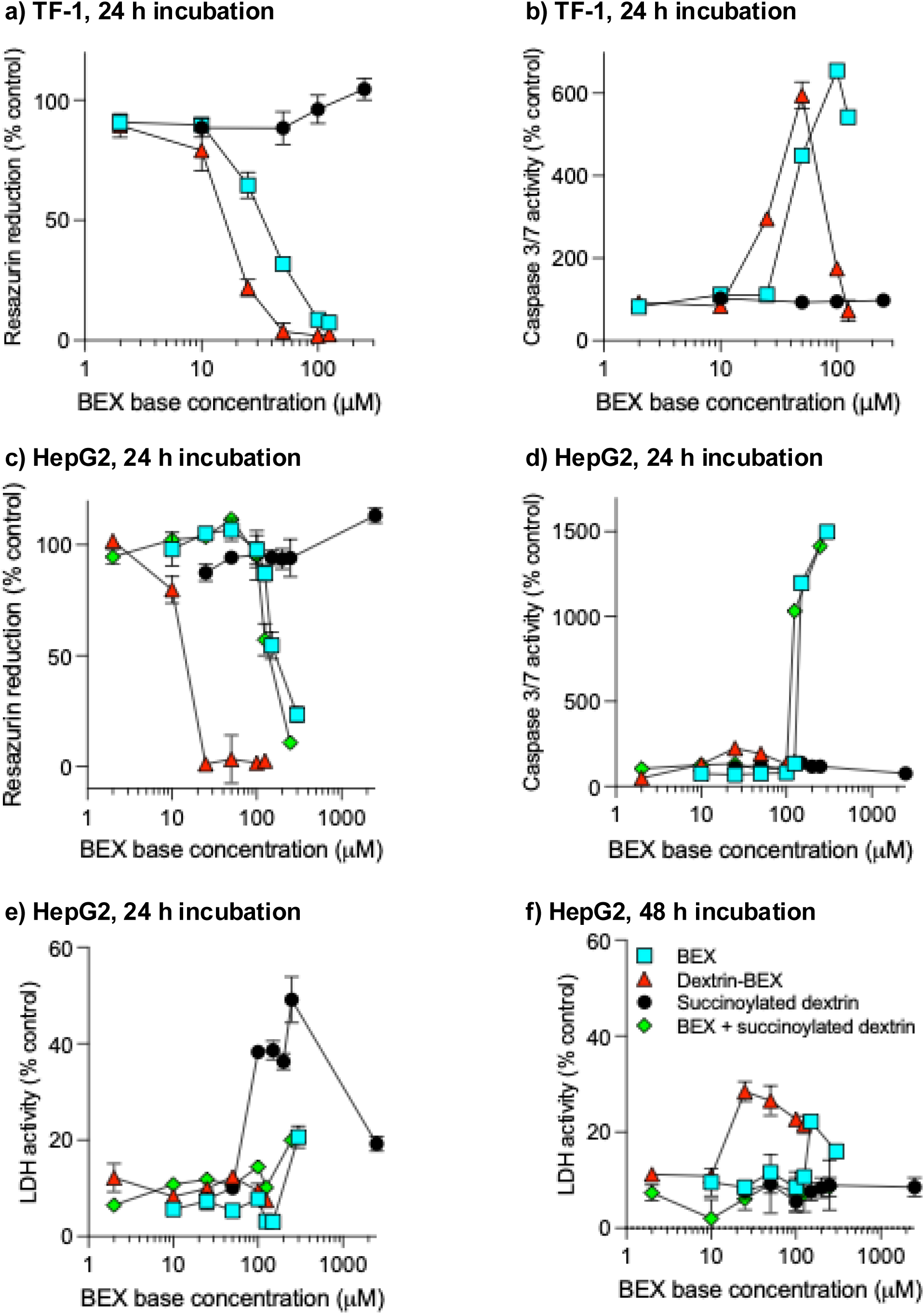
Effects of BEX formulations on cell viability, apoptosis, and membrane integrity. Multiplex analysis of resazurin reduction (metabolic activity), caspase-3/7 activation (apoptosis), and LDH release (membrane damage) in TF-1 (leukaemia) (a,b) and HepG2 (liver) (c-f) cells following 24 or 48 h exposure to BEX, dextrin-BEX conjugate, unconjugated BEX with succinoylated dextrin, or succinoylated dextrin alone at equivalent concentrations used in dextrin-BEX conjugates). Data are expressed as percentage relative to untreated controls (a–d) or lysis buffer controls (e–f) and presented as mean ± SD (n = 3). Where error bars are invisible, they are within size of data points.

**Table 2.** IC_50_ values of BEX and dextrin-BEX conjugate in leukaemia, liver and skin cell lines (24 h incubation).

| Cell line | IC<sub>50</sub> value (μM BEX base) | | |
| --- | --- | --- | --- |
|  | BEX | Dextrin-BEX conjugate<br>(fold-change vs. BEX) | Succinoylated dextrin + BEX<br>(fold-change vs. BEX) |
| MV-4-11 | 48.6 | 17.3 (2.8) | ND |
| THP-1 | 76.0 | 24.6 (3.1) | ND |
| TF-1 | 33.8 | 15.8 (2.1) | ND |
| HepG2 | 153.7 | 11.0 (14.0) | 128.3 (1.2) |
| Huh-7 | 144.7 | 17.1 (8.5) | ND |
| IHH | 133.0 | 24.9 (5.3) | 124.2 (1.1) |
| nIh | 190.7 | 19.6 (9.7), | 214.7 (0.9) |

ND, not determined

Similar patterns of cytotoxicity were observed when liver and fibroblast cell lines were treated with BEX, succinoylated dextrin and dextrin-BEX conjugates (Figure 3c-f, S2, S3, S4). When HepG2 cells were incubated with BEX for 24 h, a concentration-dependent surge in caspase 3/7-dependent apoptosis was observed that was unaltered by the addition of free succinoylated dextrin at equivalent concentrations as found in dextrin-BEX conjugate (Figure 3d). However, conjugation of succinoylated dextrin to BEX suppressed the caspase 3/7 activity surge induced by the free drug, but not to the level of the vehicle control. Succinoylated dextrin did not induce caspase 3/7 activity in any of the cell lines tested (Figure S4). Time- and concentration-dependent necrosis was observed in all cell lines, with most treatments showing no effect on LDH release at 24 h, except at the very highest concentrations (Figure S3). Succinoylated dextrin induced LDH release at polymer concentrations equivalent to >50 µM BEX base without loss of viability, which was undetectable by 48 h (Figure 3e,f). By 48 h, BEX and dextrin-BEX conjugate caused a concentration-dependent increase in LDH release that occurred at lower concentrations for the conjugate than the free drug (25 vs. 150 µM BEX base) (Figure 3f).

### Intracellular triglycerides

Treatment of immortalised and tumour liver cell lines (HepG2, Huh7 and IHH) with dextrin-BEX conjugate for up to 48 h did not cause any significant intracellular triglyceride accumulation (Figure 4). However, reduced triglyceride levels were observed when IHH cells (immortalised human hepatocytes) were treated with higher concentrations of BEX (50-100 µM, IC_50_ = 133.0 µM).

**Figure 4.**
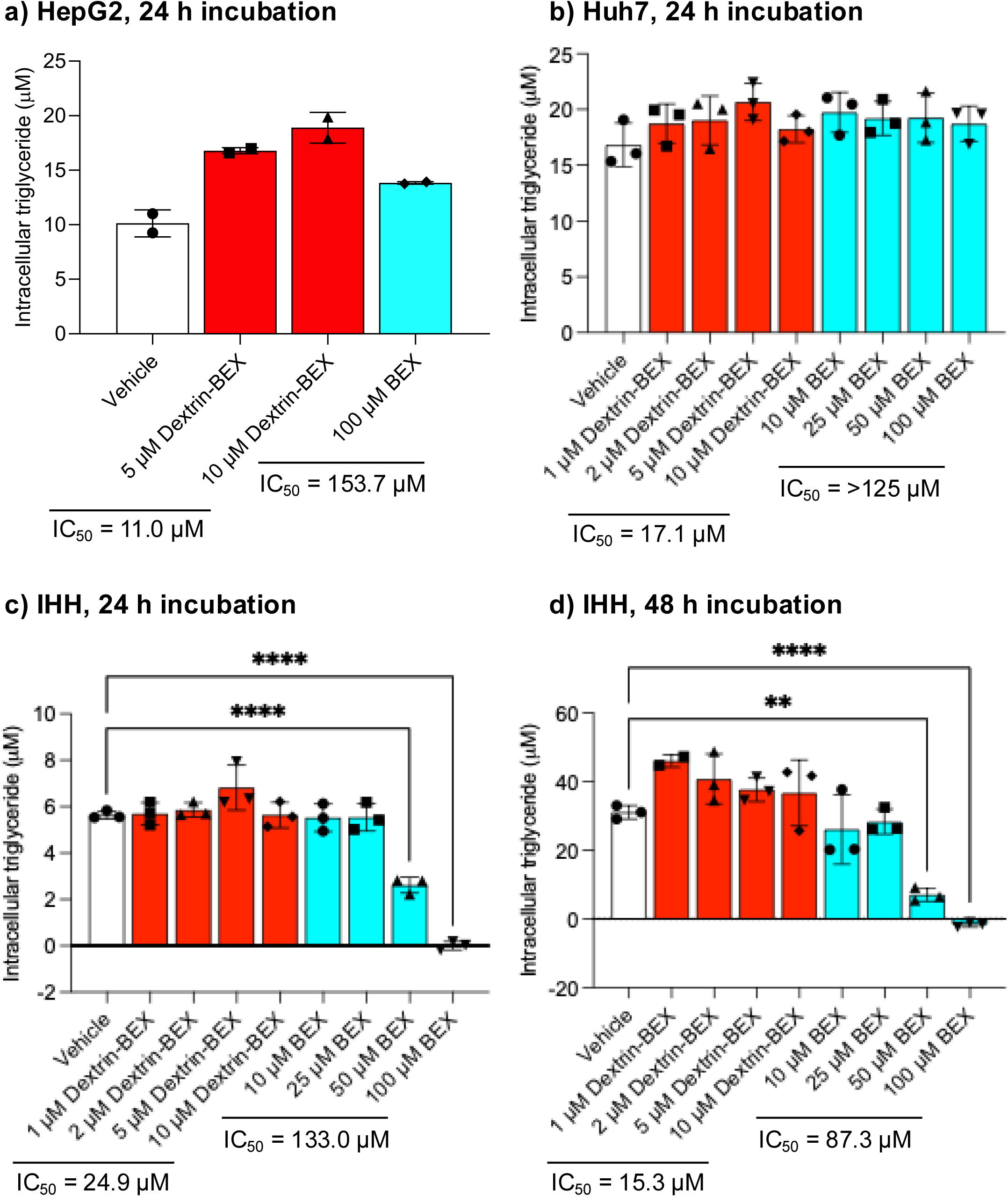
Intracellular triglyceride accumulation following treatment with BEX or dextrin-BEX. Triglyceride levels in (a) HepG2, (b) Huh7, and (c,d) IHH cells after 24 or 48 h exposure to increasing concentrations of BEX or dextrin-BEX conjugate. IC_₅₀_ values are indicated where applicable. Data are presented as mean ± SD (n = 2-3). Statistical significance where n=3: **p < 0.01, ****p < 0.0001 versus vehicle; ns, p > 0.05. Where significance is not shown, *p*>0.05 (not significant).

### *In vivo* mouse model

None of the treatments caused significant weight loss in mice after 14 days of treatment; indeed, mice in all groups gained weight during the 14-day study, with no significant differences between treatments (Figure 5a). In contrast, treatment with BEX or the dextrin-BEX conjugate for 14 days increased liver weight by 129% and 117.3%, respectively, compared with saline treatment (Figure 5b,c). Although free BEX caused a significant increase in liver weight compared with saline, the dextrin-BEX conjugate’s effect was not significant. None of the treatments significantly altered serum AST activity, serum and liver triglyceride levels or liver cytokine levels (Figure 6).

**Figure 5.**
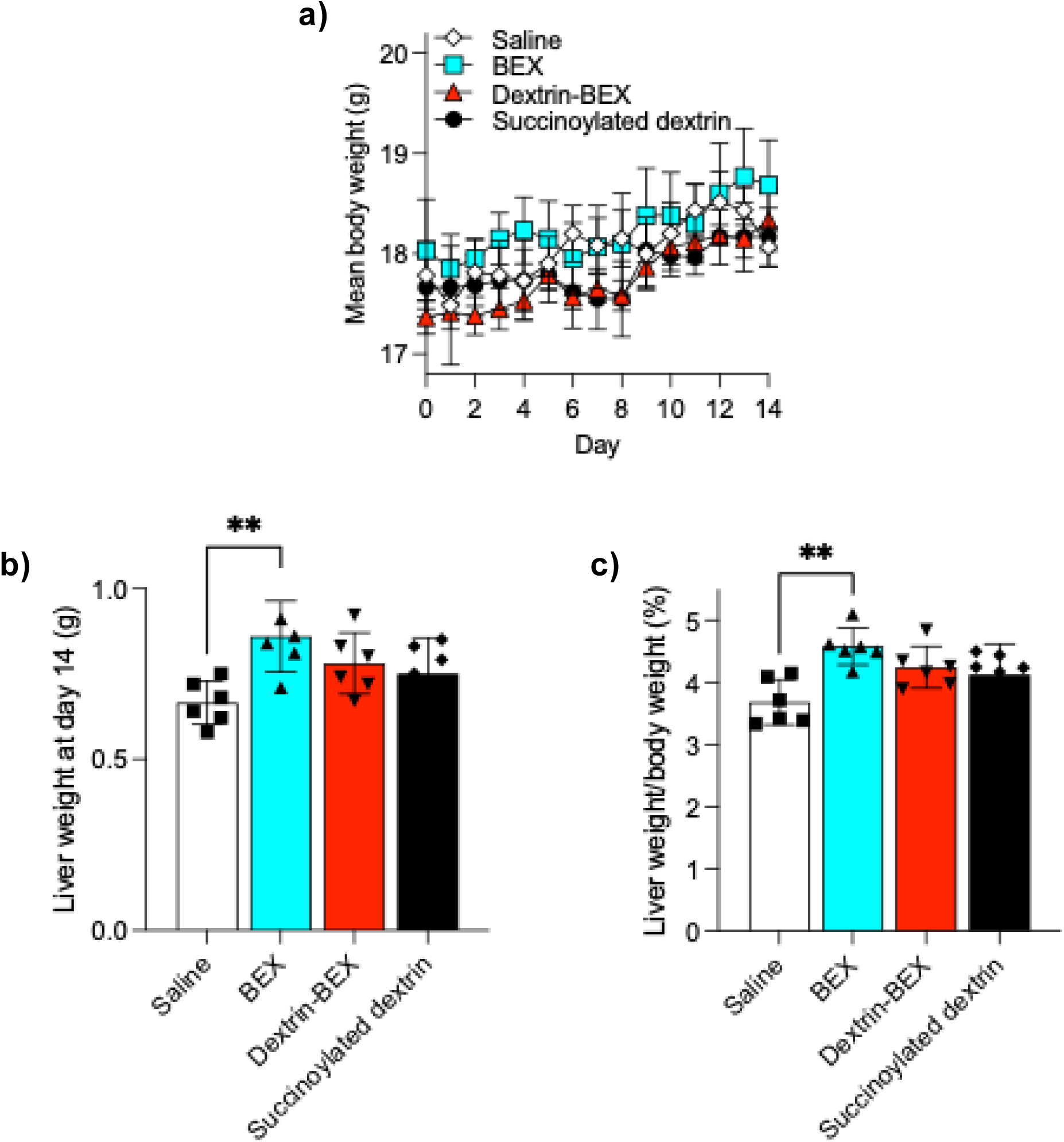
Effects of treatment on body and liver weight in vivo. Panel a) shows mean body weight over14 days following daily administration of saline (control), BEX, dextrin-BEX conjugate or succinoylated dextrin. Panel b) shows liver weight at day 14 and panel c) shows liver-to-body weight ratio at day 14. Data are presented as mean ± SEM (n = 6). **p < 0.01 versus saline; ns, p > 0.05. Where significance is not shown, *p*>0.05 (not significant).

**Figure 6.**
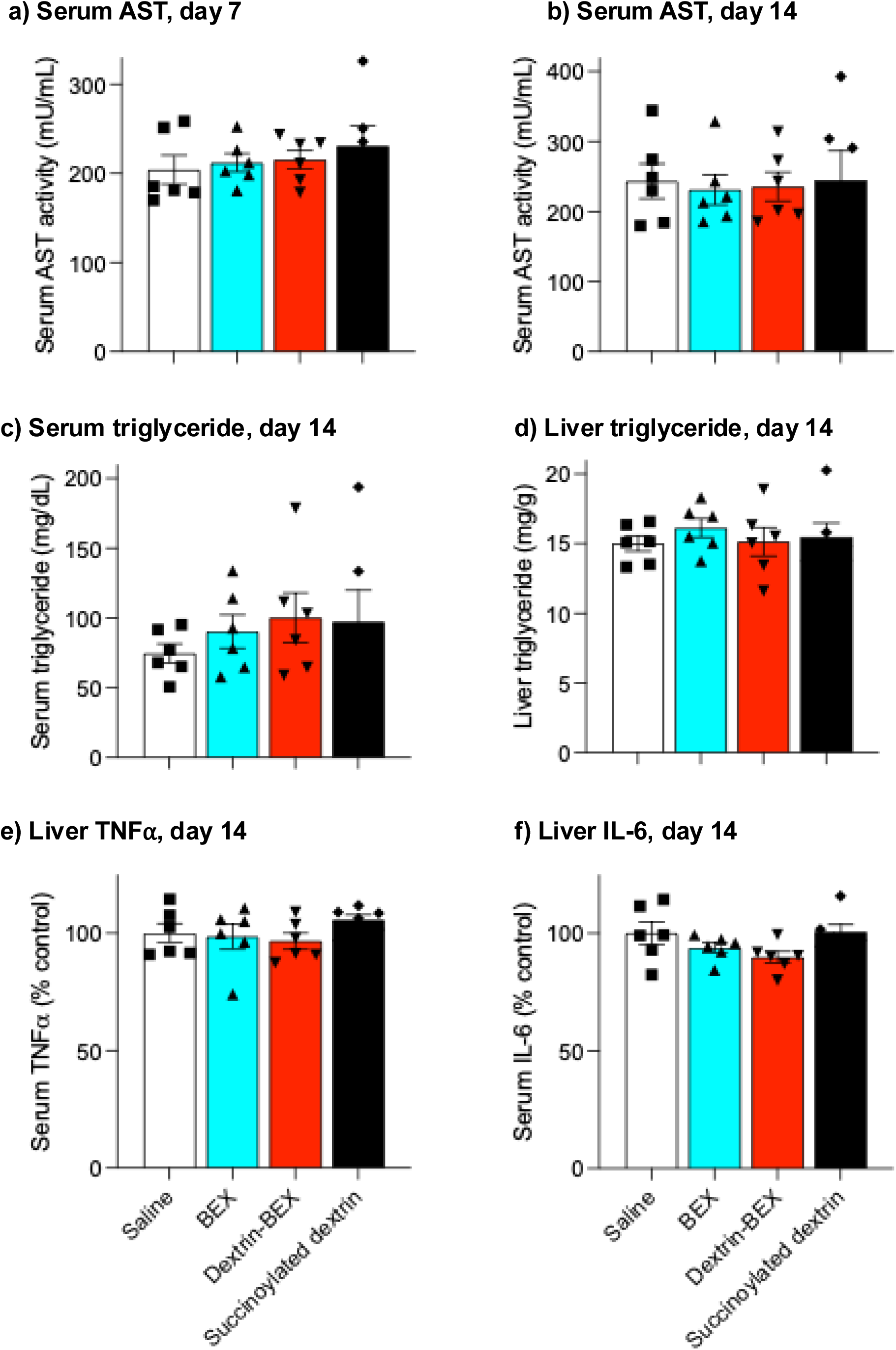
Biochemical markers of liver function, lipid metabolism, and inflammation in treated mice. Serum aspartate aminotransferase (AST) at (a) day 7 and (b) day 14; (c) serum triglycerides and (d) liver triglycerides at day 14; hepatic (e) TNF-_α_ and (f) IL-6 levels at day 14 following daily treatment with saline, BEX, dextrin-BEX conjugate, or succinoylated dextrin. Data are presented as mean ± SEM (n = 6). Where significance is not shown, *p*>0.05 (not significant).

BEX levels in serum and liver tissue harvested from mice 12 h after the final dose of BEX or dextrin-BEX conjugate were below the quantification limit (Figure S6, Table S1).

## Discussion

Bexarotene has demonstrated clinical efficacy in CTCL and considerable therapeutic potential in a range of diseases. However, its broader clinical translation remains limited by poor aqueous solubility, low bioavailability and a high incidence of dose-limiting adverse effects, most notably hyperlipidaemia. As a result, there has been substantial interest in developing formulation strategies to improve its therapeutic index. This study sought to determine whether dextrin conjugation could address these limitations by improving solubility, modulating drug disposition and reducing adverse effects.

Here, BEX was conjugated to dextrin via an ester bond using BEX’s carboxyl group, meaning that incorporation of a reactive group to dextrin was not necessary. However, succinoylation of dextrin has been shown to delay amylase-mediated polymer degradation,^32, 33^ thereby extending the release of BEX. Thus, we chose a high molecular weight dextrin with 30mol% succinoylation for attachment to BEX, as our previous studies showed that this would ensure prolonged drug release for at least 48 h.^34^ As expected, dextrin conjugation substantially enhanced aqueous solubility, increasing BEX solubility from <0.004 mg/mL to approximately 9.88 mg/mL. This represents several orders of magnitude improvement and exceeds the solubility gains reported for many alternative delivery systems, including micelles and co-crystals.^9, 12, 15–17^ This dramatic increase can be attributed to the amphiphilic nature of the conjugate, in which hydrophobic BEX moieties are incorporated into a hydrophilic dextrin backbone. DLS analysis indicated that this structure promotes self-assembly into nanoscale aggregates, driven by hydrophobic interactions between BEX residues, resulting in a stable aqueous dispersion. These physicochemical changes are likely responsible for the observed increase in *in vitro* potency, with IC50 values reduced by up to 14-fold across multiple cell lines. The formation of nanoscale aggregates may facilitate cellular uptake via endocytic pathways and increase the drug’s intracellular availability. However, unlike targeted systems such as HA-BEX conjugates or folate-functionalised nanoparticles, where enhanced efficacy is linked to receptor-mediated delivery,^10, 13, 14, 16^ the increased potency observed for dextrin-BEX appears to be driven primarily by improved solubility and non-specific uptake rather than selective targeting.

Importantly, the same aggregation behaviour that enhances solubility and uptake may also limit drug release. Dense polymeric assemblies can restrict access of ester linkages to enzymatic cleavage, reducing the rate at which free BEX is released. This may explain the absence of detectable drug levels in serum and liver *in vivo,* suggesting either slow release, rapid clearance of degradation products or limitations in analytical sensitivity. In contrast, other delivery systems, such as nanocrystals and co-crystals, have demonstrated measurable improvements in pharmacokinetics, including increased bioavailability and systemic exposure.^9, 16^

A major limitation identified in this study is the lack of selectivity of dextrin-BEX conjugates. Although cytotoxicity was enhanced in cancer cell lines, similar effects were observed in normal fibroblast and liver cells. This non-selective toxicity reflects both the biology of BEX and the properties of the delivery system. RXR receptors are ubiquitously expressed across normal and malignant tissues, meaning that increased intracellular delivery of BEX can amplify pharmacological effects in both. Furthermore, many downstream pathways regulated by RXR, including lipid metabolism, apoptosis and inflammation, are fundamental to normal cellular function and are not cancer-specific. Additional factors may contribute to this lack of selectivity. Drugs that disrupt core cellular processes, such as transcriptional regulation or metabolic homeostasis, can affect both normal and malignant cells. Similarly, cells with high metabolic activity or turnover may be particularly susceptible to such agents. In the absence of a targeting mechanism, dextrin-BEX conjugate uptake is likely mediated by non-specific processes, such as endocytosis of polymeric aggregates, further increasing off-target effects. This contrasts with targeted delivery systems, such as HA-based or ligand-functionalised nanoparticles (e.g., folate-modified nanoparticles), which exploit receptor overexpression to achieve preferential uptake in tumour cells and reduce toxicity in normal tissues.^10, 14^

The central aim of this study was to determine whether dextrin conjugation could mitigate BEX-induced lipid dysregulation. Polymer conjugation is a well-known and widely used technique to reduce drug side effects.^18^ Indeed, in our recent studies, conjugation of dextrin to the antibiotic colistin dramatically diminished proximal tubular injury and colistin renal accumulation in a murine model of acute kidney injury.^21^ Therefore, we hypothesised that attaching dextrin to BEX could reduce the drug-induced lipid abnormalities. However, neither the *in vitro* nor the *in vivo* data support this hypothesis. In hepatic cell lines, the dextrin-BEX conjugate did not significantly alter intracellular triglyceride levels compared with controls, suggesting that conjugation does not prevent BEX-induced modulation of lipid metabolism at the cellular level. Although intracellular triglyceride levels dropped at higher free BEX concentrations, this likely results from cytotoxic effects such as membrane disruption and early stages of cell death, rather than a genuine impact on lipid metabolism.

*In vivo*, both free and dextrin-conjugated BEX increased liver weight, consistent with hepatic steatosis reported in previous BEX studies.^35, 36^ Importantly, this increase was not accompanied by elevated AST levels or changes in inflammatory cytokines, suggesting that while steatosis occurred, there was no evidence of acute hepatocellular injury. This distinction is important, as it indicates that BEX-induced lipid accumulation may occur independently of overt liver damage. The inability of dextrin conjugation to mitigate liver enlargement or triglyceride accumulation suggests that the underlying mechanism of dyslipidaemia is not addressed by this formulation approach. This finding is consistent with observations from other BEX delivery systems. Across a broad range of strategies, including nanocrystals, co-crystals, micelles, liposomes and polymer conjugates, there is currently no convincing evidence that formulation approaches alone can prevent BEX-induced hypertriglyceridaemia. Most systems have been designed to improve solubility, enhance bioavailability or enable tissue targeting, rather than to address systemic metabolic effects. This suggests that hyperlipidaemia is an inherent pharmacodynamic consequence of RXR activation, likely mediated through alterations in lipid homeostasis pathways, and therefore represents a fundamental limitation of the drug rather than a purely formulation-dependent issue.

When positioned within the wider landscape of delivery strategies, dextrin-BEX conjugate exhibits a distinct profile. It achieves exceptional solubility enhancement and improves *in vitro* potency, placing it among the most effective systems in terms of physicochemical performance. However, it lacks the pharmacokinetic advantages of systems such as nanocrystals and co-crystals and does not provide the selectivity associated with targeted platforms.^9, 10, 14, 16^ Most importantly, it does not address the central clinical limitation of BEX, namely metabolic toxicity. These findings suggest that improving the therapeutic index of BEX will require more sophisticated, multi-functional delivery systems that combine solubility enhancement with active targeting and controlled drug release. Furthermore, mitigation of dyslipidaemia may require complementary pharmacological strategies, such as lipid-lowering therapies or the development of RXR modulators with reduced metabolic liability.

## Conclusion

In conclusion, while dextrin conjugation is an effective strategy for improving the solubility and cellular potency of BEX, it does not improve its safety profile or mitigate its key adverse effects *in vivo*. These results, together with evidence from other delivery systems, indicate that overcoming BEX-induced dyslipidaemia remains a fundamental challenge and a critical barrier to its broader clinical application.

## Author contributions

ELF and SR conceived the project. ELF, SR, TRH and YZ designed the study and SR, CP, TA, GB, TA, CA, TR, TRH and ELF performed the experiments. ELF, SR, TRH and YZ acquired the funding. ELF took the lead in writing the manuscript. All authors provided critical feedback and helped shape the research, analysis and manuscript.

## Data availability

Information on the data underpinning the results presented here, including how to access them, can be found in the Cardiff University data catalogue at (TBC)

## Acknowledgements

This research was funded in whole, or in part, by the Wellcome Trust (204824/Z/16/Z). This research was also funded by Cardiff University. For the purpose of open access, the author has applied a CC BY public copyright license to any Author Accepted Manuscript version arising from this submission. The funders had no role in the design of the study; in the collection, analyses, or interpretation of data; in the writing of the manuscript, or in the decision to publish the results. The authors thank Mrs Nicola Beynon for inspiring this research. We also acknowledge the technical cell culture support provided by Dr Sarah Youde and Dr Maria Stack. During preparation, this manuscript contributed to a user testing study for the <u>RIVER recommendations</u> and was reviewed by members of the <u>RIVER recommendations working group</u>’. This paper has been uploaded to bioRxiv as a preprint:

