## Supplementary files for "Development of a nanomedicine approach to improve the therapeutic index of bexarotene"

**Table S1.** Characteristics of cell lines used in this study

| Cell line | Source | Patient | Pathology | Culture medium[1] | RXR expression[2] |
|---|---|---|---|---|---|
| MV-4-11 | Peripheral blood | Caucasian male, 10 Y | Biphenotypic B myelomonocytic leukaemia | IMDM with GlutaMAX™, 10% v/v heat-inactivated (HI)-FBS | RXRA: 5.23<br>RXRB: 4.76<br>RXRG: 0 |
| THP-1 | Peripheral blood | Japanese male, 1Y | Acute monocytic leukaemia | RPMI 1640 withGlutaMAX™, 10% HI-FBS | RXRA: 5.93<br>RXRB: 6.08<br>RXRG: 0 |
| TF-1 | Bone marrow | Japanese male, 35 Y | Erythroleukaemia | RPMI-1640, with GlutaMAX™,10% v/v HI-FBS, 2 ng/mL rhGM-CSF | RXRA: 3.61<br>RXRB: 4.70<br>RXRG: 0 |
| HepG2 | Liver | Caucasian male, 15 Y | Hepatoblastoma | DMEM high glucose, 10% v/v HI-FBS, GlutaMAX | RXRA: 4.36<br>RXRB: 4.53<br>RXRG: 0 |
| Huh-7 | Liver | Japanese male, 57 Y | Hepatocellular carcinoma | DMEM high glucose, 10% v/v HI-FBS, GlutaMAX | RXRA: 5.43<br>RXRB: 4.25<br>RXRG: 0.01 |
| IHH | Liver | Male, 59 Y | Hepatocyte | Williams' medium E with 20 mU/mL insulin +10% v/v HI-FBS | ND |
| nIh | Skin | Female, 75 Y | Fibroblast | DMEM high glucose 10% v/v HI-FBS + 1x amino acid supplement | ND |

RXR expression data (RXRA, RXRB, RXRG) were obtained from DepMap (DepMap, Broad (2024). DepMap 24Q4 Public. Figshare+. Dataset. https://doi.org/10.25452/figshare.plus.27993248.v1]) and are reported as $\log_2$(TPM + 1). Culture media did not contain antibiotics or antimycotics. ND, not determined.

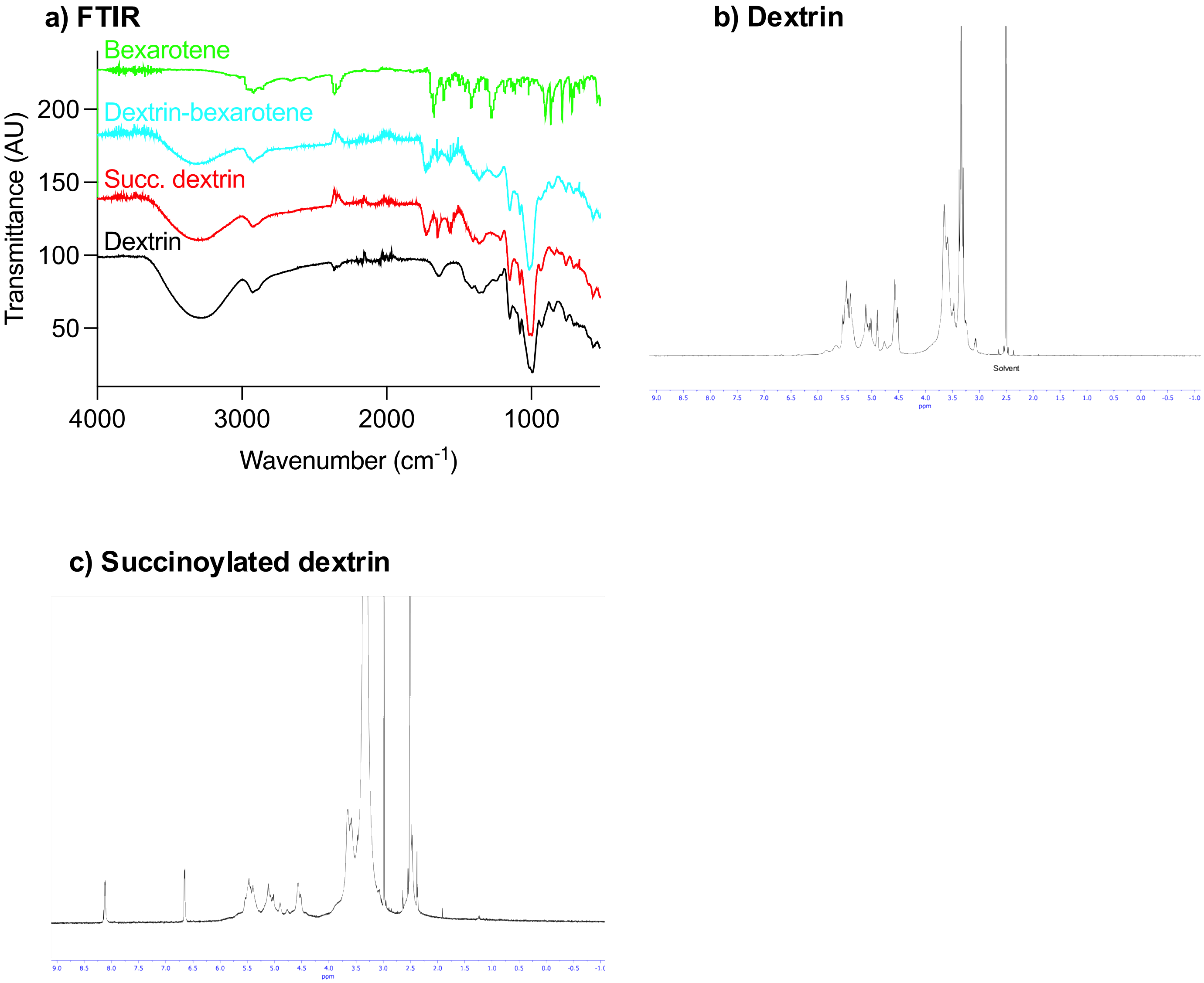


**Figure S1.** Physicochemical characterisation of starting materials and intermediates. Panel (a) shows FTIR spectra of dextrin, succinoylated dextrin, dextrin-BEX and BEX. Panels (b-d) show $^{1}$H-NMR spectra of dextrin and succinoylated dextrin.

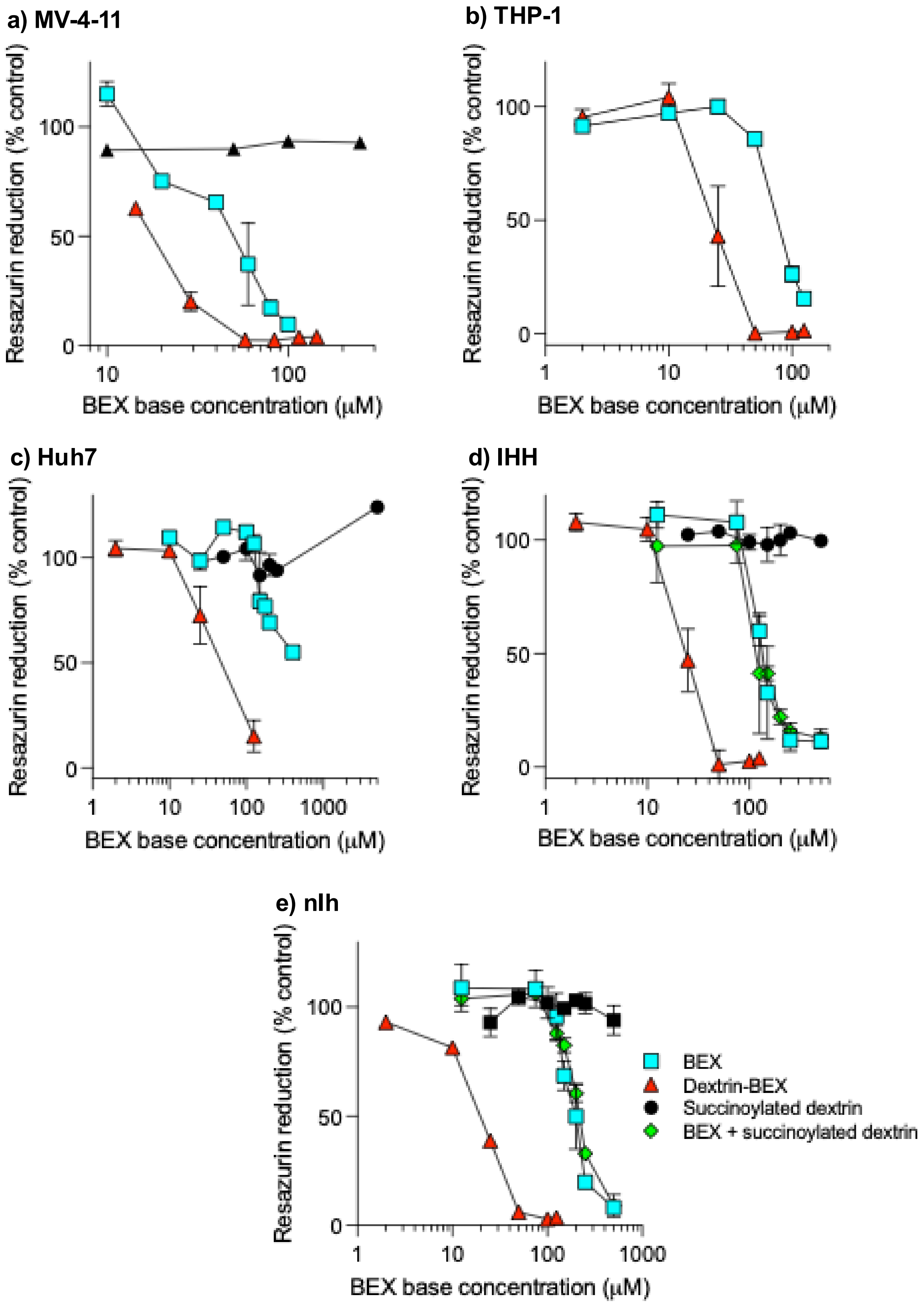


**Figure S2.** Metabolic activity of additional cell lines following treatment. Resazurin reduction (CellTiter Blue) measured in (a) MV-4-11, (b) THP-1, (c) Huh7, (d) IHH, and (e) nIh cells after 24 h exposure to BEX, dextrin-BEX conjugate, unconjugated BEX with succinoylated dextrin, or succinoylated dextrin alone at equivalent concentrations. Data are expressed as percentage of vehicle control (0.5% DMSO) and presented as mean ± SD (n = 3). Where error bars are invisible, they are within size of data points.

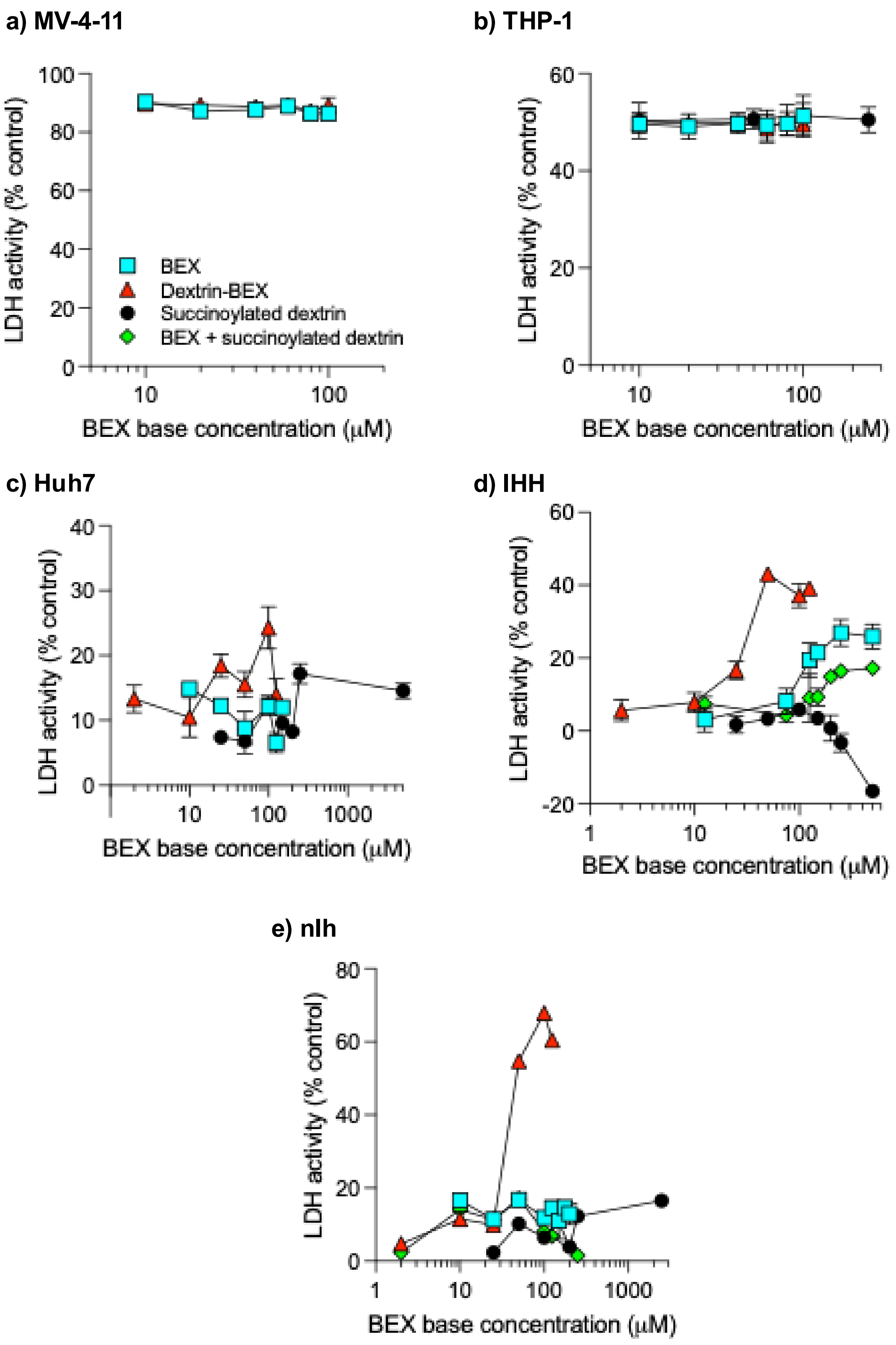


**Figure S3.** Membrane integrity (necrosis) in additional cell lines. LDH release measured in (a) MV-4-11, (b) THP-1, (c) Huh7, (d) IHH, and (e) nIh cells after 24 h treatment with BEX, dextrin-BEX conjugate, unconjugated BEX with succinoylated dextrin, or succinoylated dextrin alone. Data are expressed as percentage of lysis buffer control and presented as mean ± SD (n = 3). Where error bars are invisible, they are within size of data points.

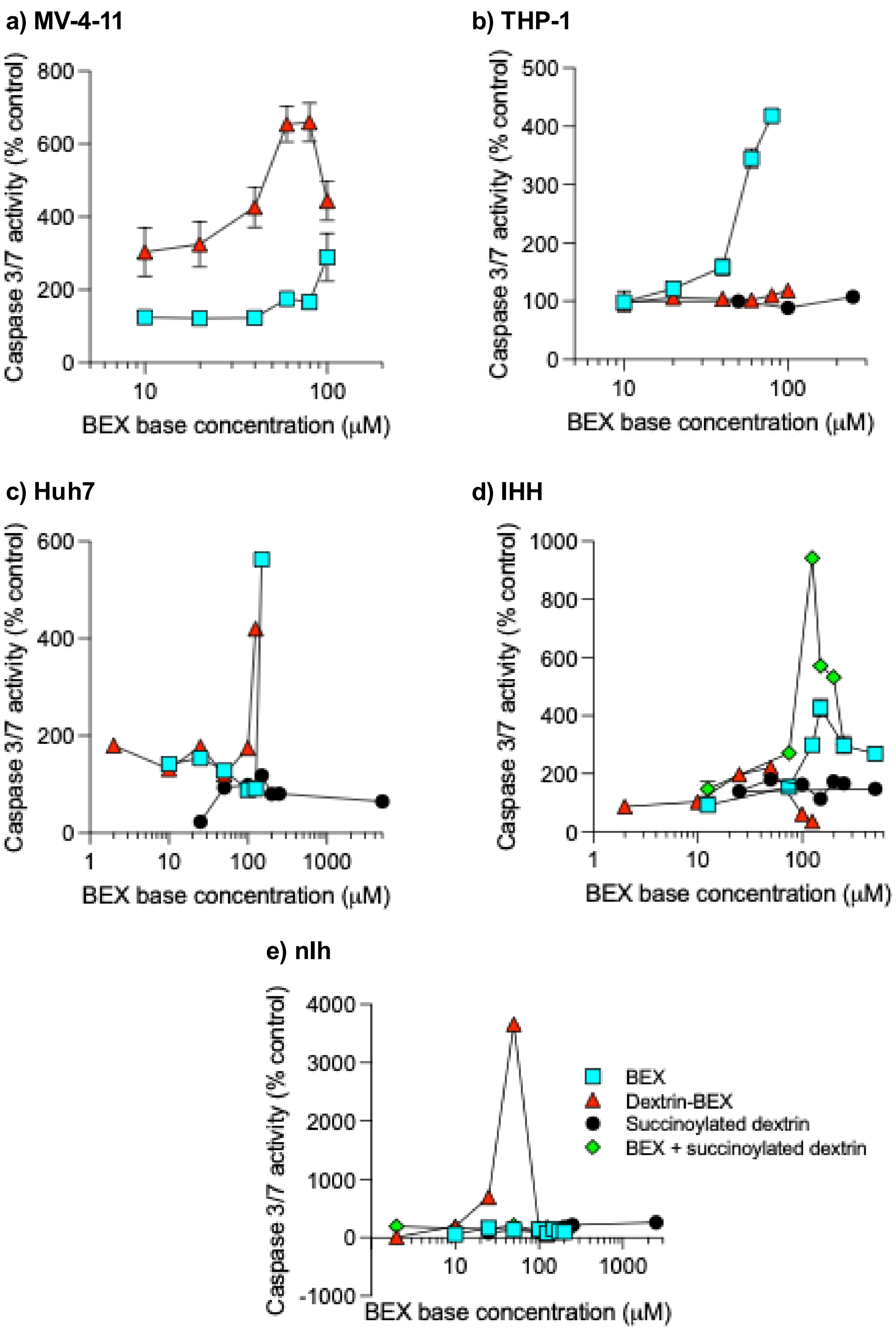


**Figure S4.** Apoptosis induction in additional cell lines. Caspase-3/7 activity measured in (a) MV-4-11, (b) THP-1, (c) Huh7, (d) IHH, and (e) nIh cells after 24 h exposure to BEX, Dex-BEX conjugate, unconjugated BEX with succinoylated dextrin, or succinoylated dextrin alone. Data are expressed as percentage of vehicle control (0.5% DMSO) and presented as mean ± SD (n = 3). Where error bars are invisible, they are within size of data points.

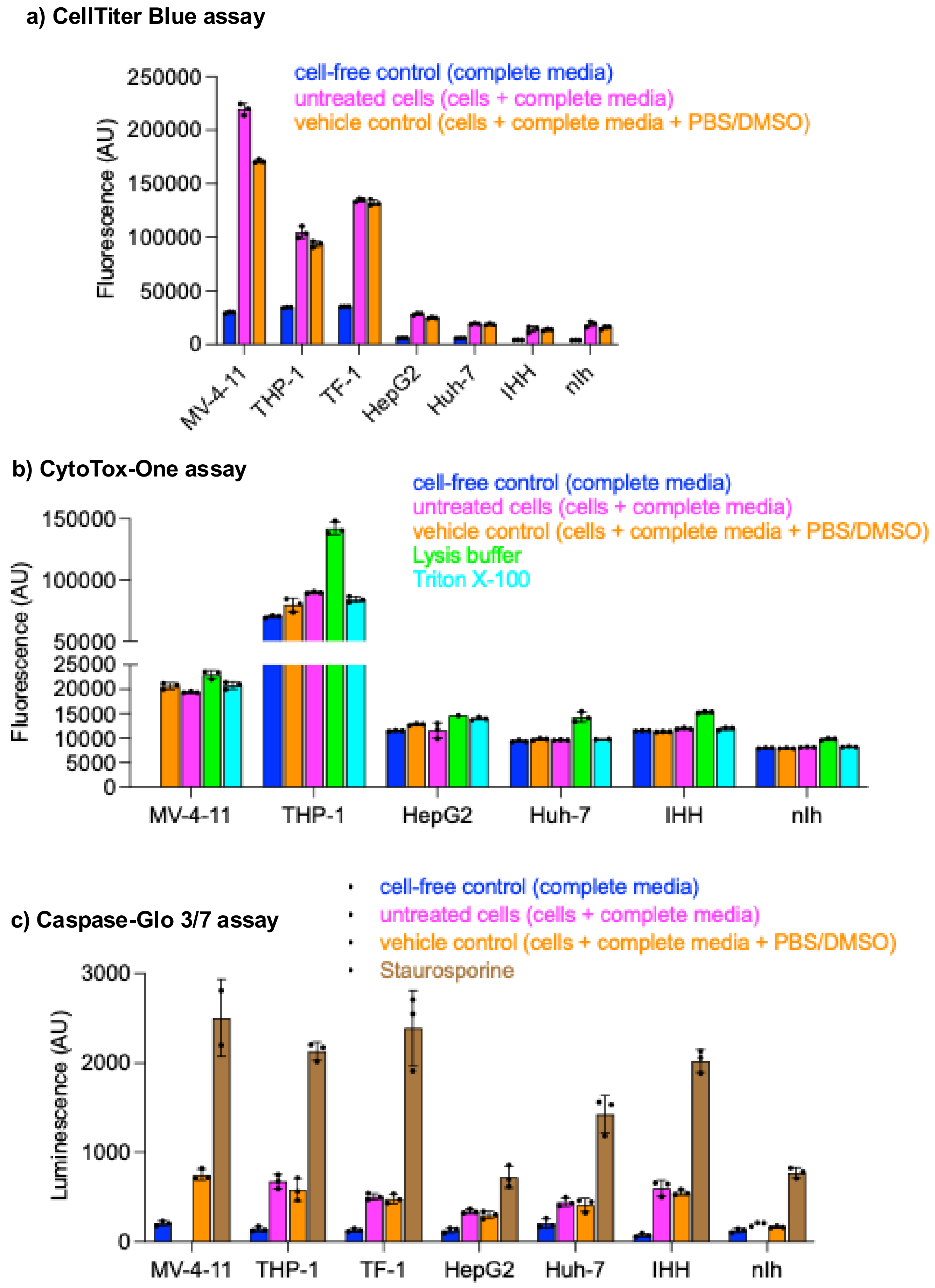


**Figure S5.** Summary of assay control data for multiplex cytotoxicity assays. Panel (a) shows CellTiter Blue (metabolic activity), (b) CytoTox-One (LDH release), and (c) Caspase-Glo® 3/7 (apoptosis) control signals. Data are presented as fluorescence (a,b) or luminescence (c) values (mean ± SD, n = 3).

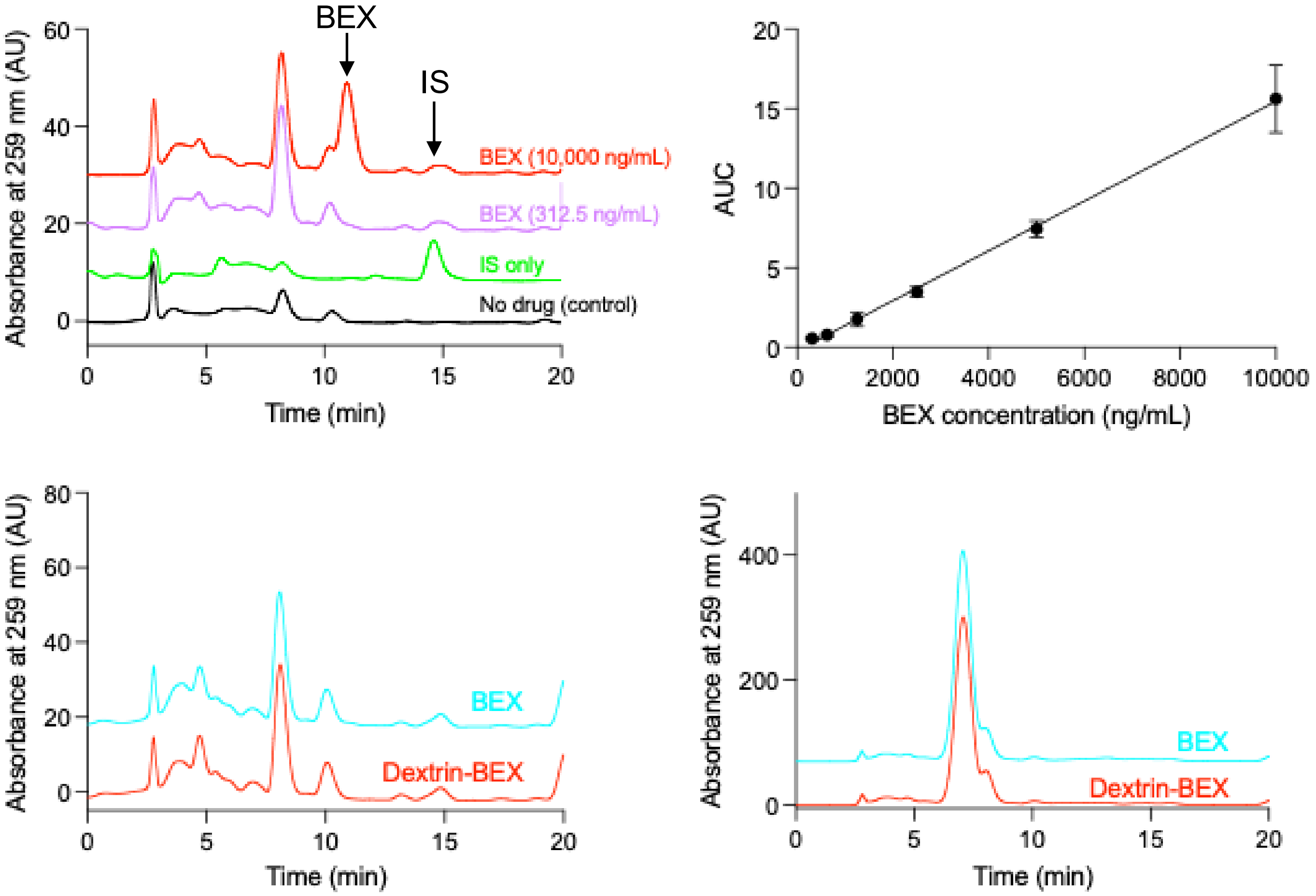


**Figure S6.** Representative HPLC-UV chromatograms for quantification of BEX in biological samples. Panel a) shows chromatograms recorded at λ = 259 nm of FBS spiked with BEX at low and high BEX concentration, and panel b) shows a BEX calibration curve. Representative c) serum and d) liver samples collected 12 h after the final dose of BEX or dextrin-BEX conjugate are also shown.

**Table S1: L**imits of detection, quantification and regression for the analysis of BEX concentration

| | |
|---|---|
| Linearity range | 312.5 – 10,000 mg/mL |
| Correlation coefficient ($r^2$) value | 0.9787 |
| Regression equation | Y = 0.001566*X - 0.1804 |
| LOD (Limit of detection) $= \frac{3.3 \times \sigma}{m}$ | 1828.3 |
| LOQ (Limit of quantification) $= \frac{10 \times \sigma}{m}$ | 5540.2 |

where $\sigma$ = standard deviation and $m$ = slope of calibration curve
